# Structural Modeling of Chromatin Integrates Genome Features and Reveals Chromosome Folding Principle

**DOI:** 10.1101/085167

**Authors:** Wen Jun Xie, Luming Meng, Sirui Liu, Ling Zhang, Xiaoxia Cai, Yi Qin Gao

**Author notes:** Correspondence (Y.Q.G).

## Abstract

How chromosomes fold into 3D structures and how genome functions are affected or even controlled by their spatial organization remain challenging questions. Hi-C experiment has provided important structural insights for chromosome, and Hi-C data are used here to construct the 3D chromatin structure which are characterized by two spatially segregated chromatin compartments A and B. By mapping a plethora of genome features onto the constructed 3D chromatin model, we show vividly the close connection between genome properties and the spatial organization of chromatin. We are able to dissect the whole chromatin into two types of chromatin domains which have clearly different Hi-C contact patterns as well as different sizes of chromatin loops. The two chromatin types can be respectively regarded as the basic units of chromatin compartments A and B, and also spatially segregate from each other as the two chromatin compartments. Therefore, the chromatin loops segregate in the space according to their sizes, suggesting the excluded volume or entropic effect in chromatin compartmentalization as well as chromosome positioning. Taken together, these results provide clues to the folding principles of chromosomes, their spatial organization, and the resulted clustering of many genome features in the 3D space.

## Introduction

Eukaryotic genomes in the interphase are tightly packaged within cell nuclei, fold into compact 3D structures, and form chromosome territories^1,2^. The spatial organization of chromatin plays an essential role in various genome functions, including gene expression, DNA replication and cell mitosis^3-7^.

Hi-C, a chromosome-conformation-capture based method, has provided profound structural insights for chromosome. For example, these data suggested that chromatin fiber is spatially organized into two tissue-specific compartments (A/B compartments) in the megabase scale or tissue-invariant topologically associating domains (TADs) in the kilobase scale^8,9^. In Hi-C experiments, the relative frequencies of spatial contacts between genomic loci from a cell population and homologs are measured. Recently, the two chromatin compartments in single chromosomes are also confirmed by fluorescence in situ hybridization (FISH) which can directly image the 3D chromatin^10^. In addition, identification of chromatin loops from Hi-C data helps to explain the role of distal genomic interactions in controlling gene expression^11^.

Modeling chromatin structure using experimental Hi-C data is also indispensable in our understanding of genome properties and is expected to play increasingly important roles^12,13^. In one seminal study, Lieberman-Aiden and coworkers demonstrated that the fractal instead of the equilibrium globule is consistent with the Hi-C data at the megabase scale^8^, whereas, the loop-extrusion model was shown to explain better the Hi-C data in their later study^14,15^. Giorgetti *et al*. modeled the X inactivation center in human X chromosome and proposed that the fluctuations in chromosome conformation are coupled with transcription^16^. Zhang and Wolynes simulated the chromosome topology which was found to be largely free of knots^17^.

Chromatin compartments have been demonstrated to relate with a variety of genome features, including GC-content, replication timing, DNase I hypersensitivity and histone marks^8,12,18^. All these genome features were previously found to colocalize on a linear map, but their spatial distribution is still largely unknown. Due to the 3D nature of chromatin, the relation between genome features and 3D chromatin might provide a clear clue of how the colocalization on a linear map happens. Besides, there are still many gaps in the way to a comprehensive understanding of chromosomes in the highly crowding nucleus. For example, the mechanism of chromatin compartmentalization and chromosome positioning which are important regulatory factors in gene expression remains unknown.

Here, we explore the spatial organization of human chromatin by physical modeling utilizing experimental Hi-C data. The modeled 3D chromatin structure presents an integrated view of a large variety of genome features and provides important clues to the chromosome folding principle. We map a large number of genome features onto the modeled structure of an individual chromosome, revealing remarkable relations between them. Although lamina-associated domain and several histone marks have been mapped onto the modeled chromatin structure in previous studies^19,20^, a more comprehensive analysis of the genome features in the 3D space is in need to understanding the role of 3D chromatin. The nearly quantitative consistency between various biological data and the modeled chromatin structure not only reveals their relation but also validates our modeling. Furthermore, inspired by the different Hi-C patterns between partially methylated domain (PMD) and non-PMD (genomic region that isn’t PMD), we divided an individual chromosome into two types of chromatin domains which form the basis of higher-level chromatin compartments. The two domains have different densities of GC-content and CCCTC-binding factor (CTCF) binding site, and sizes of chromatin loops or TADs. Thus, chromatin compartmentalization can be viewed as a result of the segregation of chromatin loops or TADs according to their sizes, highlighting the importance of entropy in driving chromatin compartmentalization. We also found chromosomes position in the nucleus following this entropy-driven mechanism.

## Results

### Structural modeling of human chromosome

Here we developed a restraint-based modeling strategy to build chromatin structure. We first coarse-grained the genome structure with a bead-spring model, each bead representing a 50-kb genomic region. The resolution was chosen based on available experimental data and to save computational cost. The contact frequencies between loci in experimental Hi-C data were correlated with their spatial distances and thus we converted the Hi-C map into a set of restraints, which were then satisfied by numerical optimization using molecular dynamics simulations. We obtained a population of models from optimizing an ensemble of randomly generated initial chromosome configurations. The modeling details can be found in the Methods part of this paper. The experimental Hi-C data for IMR90 used here are obtained from Rao *et al*.^11^.

The consistency between our modeling and Hi-C experiment is evaluated based on the distribution of the modeled and restraint distance ratios of all the pairs of beads. For the conformation ensemble, 86.4% of all the simulated distances are within 0.5 to 2-fold of the corresponding restraint distances (Fig. S1). The comparison between contact frequencies obtained in experiments and the modeling for chromosome 1 in IMR90 is shown in Fig. 1a. The performance of the model is demonstrated at a finer scale by zooming-in the central region into Fig. 1b (or Fig. S2). The agreement between the experimental and calculated results clearly demonstrates the accuracy that our modeling strategy has achieved. Throughout the paper, we take the modeled structure of chromosome 1 of IMR90 as an example, unless otherwise stated.

**Figure 1.**
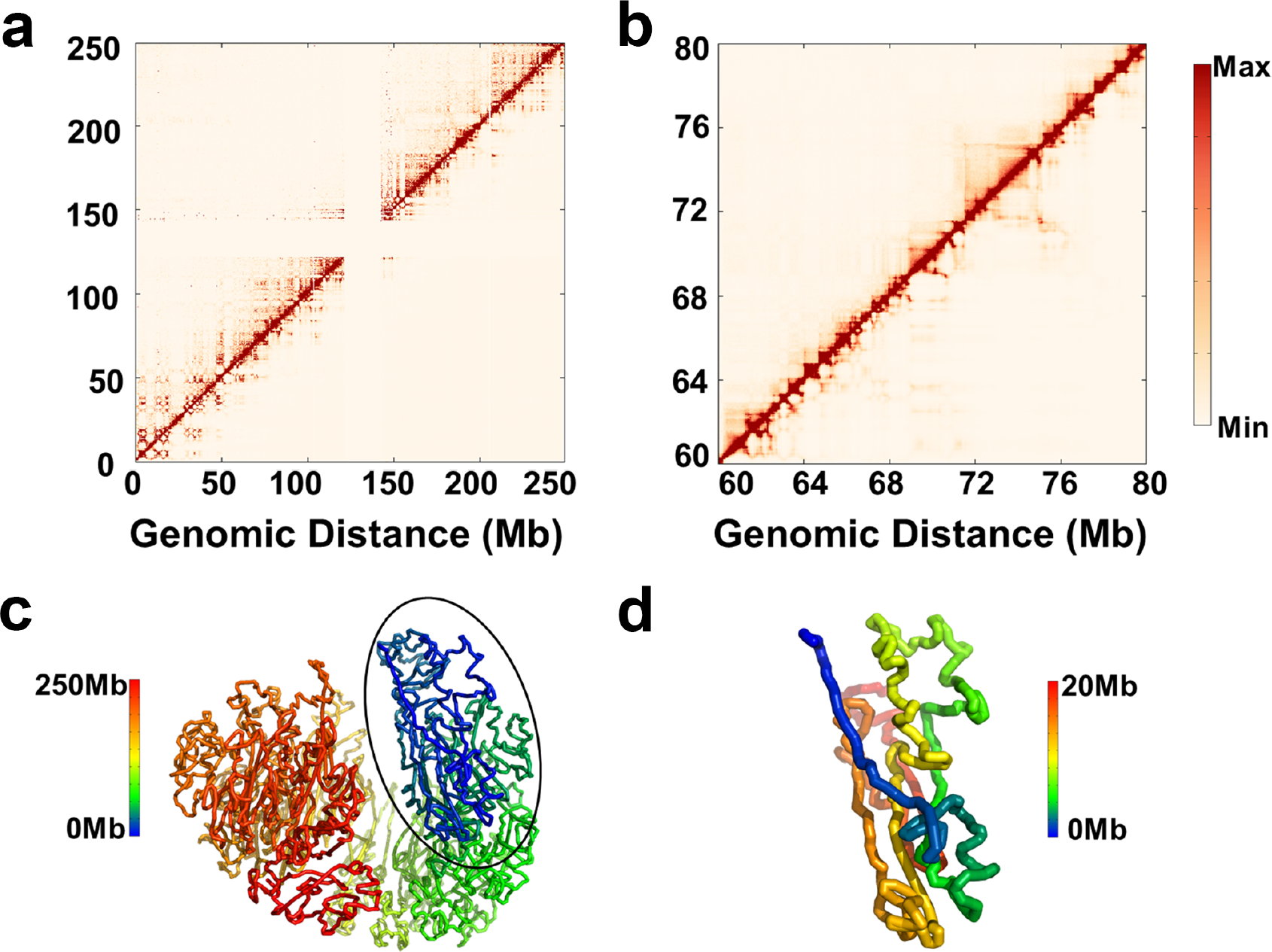
Validation of modeling and structural characterization of human chromosomes. **(a)** Comparison between experimental (upper triangle) and modeled (lower triangle) contact frequencies. **(b)** Zoomed-in version of (a) in a 20 Mb region. The consistency between the experimental and modeled contact frequencies validates the modeling strategy. **(c)** Representative modeled structure showing the packed conformation of chromosome. **(d)** Part of (c) (20 Mb) with enlarged view.

We analyzed the features of the chromosome structure ensemble. Clustering based on the root-mean-square deviation (RMSD) between any two conformations demonstrates that conformations are similar to each other and there isn’t a dominant structure in the models (Fig. S3 and SI text: Characterization of the modeled structure ensemble). But all the constructed models have a packed configuration. The cluster centroid is chosen as representative which is shown in Fig. 1c. Such a packed spherical configuration of chromosome was also found in previous modeling studies^17,21^. In Fig. S4, we also presented the structure for chromosome 18 and 19 in IMR90 which shows a packed non-spherical configuration. These packed conformations support the concept of chromosome territories that each chromosome organizes itself into a given volume in the nucleus and only contacts with its immediate neighbors^1^.

We also found that our models are largely devoid of knots (Fig. S5 and SI text: Knot invariant of the modeled structure ensembles) which has been argued to be important in chromatin folding and unfolding^8,17^. Our 3D structure also reproduces the chromatin loop identified in Hi-C data (SI text: Reproduction of chromatin loops in modeled structure)^11^. In addition, the chromatin shows interesting structural features at a finer scale which can be viewed as a collection of strands (Fig. 1d). Although it has not been explicitly mentioned before, strands can also be found from previously modeled conformation^22^. All these results strengthen the physical meaning of our modeling.

### Spatial segregation of compartments A and B

One can normalize the Hi-C contact matrix by the observed-expected method and then obtain a corresponding correlation matrix (Fig. 2a)^8^. The plaid pattern in Fig. 2a suggests a spatial segregation between two chromatin compartments.

**Figure 2.**
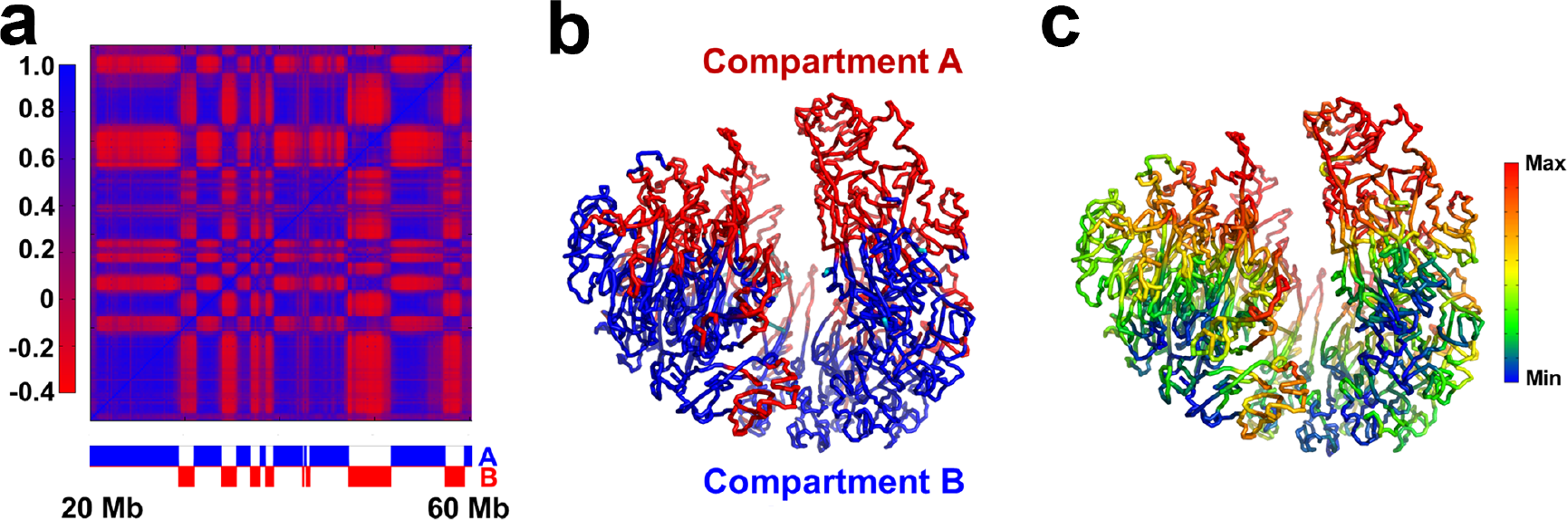
Characterization of A/B compartments. **(a)** Pearson correlation matrix. The plaid pattern suggests chromatin spatially segregates into two compartments. The assignment of each genomic region into chromatin compartments A/B is shown below. The associated degree of compartmentalization is shown in the left bar. **(b)** Spatial segregation between chromatin compartments A/B. On the representative modeled conformation, chromatin compartments A and B are shown in red and blue, respectively. **(c)** Projection of degree of compartmentalization onto representative conformation.

We then used spectral clustering to decompose human chromosome into A/B compartments (Fig. 2a and SI text: Identification of A/B compartments) and mapped their genomic regions on our model (Fig. 2b). It is obvious that the two compartments spatially segregate from each other despite that their sequences alternate along the DNA sequence (Fig. 2b). We also estimated the local density or compactness around each genomic region in our models by calculating the number of segments within a certain surrounding volume (SI text: Estimation of chromosome spatial density). The spatial density obtained from our modeling in compartment A is smaller than compartment B, implying the connection to euchromatin and heterochromatin of these two compartments, respectively.

For each genomic segment, the degree of compartmentalization is quantified as the logarithm ratio of normalized contacts with compartment A to normalized contacts with compartment B (SI text: Constructing degree of compartmentalization)^23^. The high positive degree values indicate that the genomic regions locate in the interior of compartment A whereas the low negative values correspond to the inner part of compartment B (Fig. 2c). Spatially, the interior of compartment A (red color, large positive degree values) and B (blue/green color, large negative degree values) are separated by compartment boundaries (yellow color, with degree values around zero) (Fig. 2c). The degree of compartmentalization of one genomic segment correlates with its local density (Fig. S6). In the following, we show that the definition of the degree of compartmentalization allows us to compare quantitatively the spatial packing of chromatin with genome features and reveal the biological significance of 3D chromatin structure formation.

### Spatial segregation of genome features

Here we map lamina-associated domain, DNase I hypersensitivity, RNA polymerase II, gene expression and active/repressive histone modifications to the 3D chromatin structure, due to their relation with gene activity. In addition, replication timing is also mapped because of the close relation between replication domains (RDs) with TADs. The data sources for genome features analyzed here are summarized in Table S1.

#### Lamina-associated domain

In the interphase, the nuclear lamina coating the inner nuclear membrane interacts with specific genomic regions named lamina-associated domains (LADs). LADs have long been thought to play an important role in regulating gene expression^24^. The DNA Adenine Methyltransferase Identification (DamID) technique has been used to identify LADs in a genome-wide manner^25^.

We find here that LADs can be aligned remarkably well with compartment B in the 3D space (Fig. 3a). Quantitatively, 95.5% of LAD locates in compartment B. Such a high overlapping between LAD and compartment B supports the notion that the chromatin in LADs is more compact. Given that LADs locate in the peripheral of the nucleus, our model shows that the compactness of chromatin decreases from the peripheral to central nucleus.

**Figure 3.**
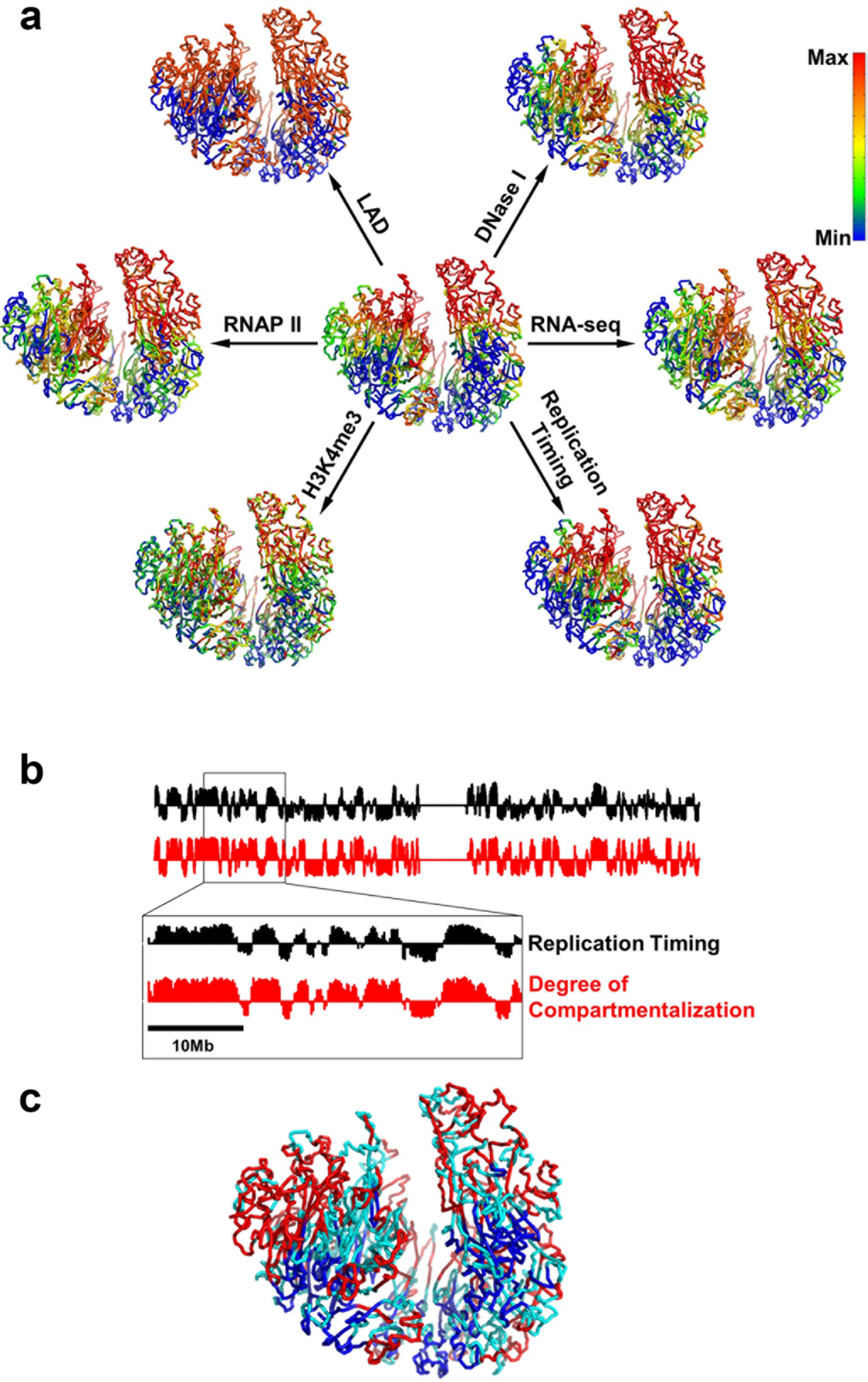
Spatial segregation of genome features. **(a)** Mapping of genome features onto the 3D chromatin structure. The degree of compartmentalization is shown in the center for comparison. Genomic features are presented as smoothed experimental data over different window sizes: 50-kb for LAD and H3K4me3, 200-kb for replication timing, 400-kb for DNase I, 2-Mb for RNA-seq and RNAP II. The maximum/minimum of color bar stands for the value of average+/−standard deviation. The red color represents deficiency of lamina-associated domain (LAD), enrichment of DNase I hypersensitive sites (DNase I), enrichment of RNA polymerase II (RNAP II), active gene expression (RNA-seq), enrichment of active histone modifications (H3K4me3) and early replication timing (Replication Timing). **(b)** Comparison between replication timing and degree of compartmentalization along the chromosome. **(c)** Spatial segregation between PMD (blue color) and non-PMD (red color). Domains with size smaller than 300 kb are not grouped into type A or B (cyan color).

#### DNase I hypersensitivity, RNA polymerase II and gene expression

By mapping the DNase I hypersensitivity data onto chromatin 3D structure (Fig. 3a), we also see that chromatin in compartment A is more accessible, which might cause the enrichment of RNA polymerase II (RNAP II) in this compartment (Fig. 3a). These results are in line with the transcriptional activation in compartment A (Fig. 3a) and highlight the importance of chromatin structure as an emerging regulator of gene expression. Such analyses are also consistent with previous analyses in that most genes in compartment B or LADs are transcriptionally inactive^26^. The relation between chromatin 3D structure and these genome properties also suggests the heterogeneous character of chromatin can play a role in regulating the diffusion of transcription factors which calls for further studies.

#### Histone modifications

We next examined the relation between chromatin structure and various histone modifications (H3K4me3, H3K27ac, H3K36me3, H3K9me3 and H3K27me3). H3K4me3 which involves in the activation of gene expression is enriched in compartment A (Fig. 3a), which is also the case for the other two active histone marks (H3K27ac, H3K36me3) (Fig. S7). Meanwhile, repressive histone marks (H3K9me3, H3K27me3) show less obvious compartmentalization (Fig. S7), revealing a major difference between repressive and active histone marks. The values of Pearson correlation between degree of compartmentalization and histone marks are 0.59 for H3K4me3, 0.61 for H3K27ac, 0.61 for H3K36me3, -0.25 for H3K9me3 and 0.29 for H3K27me3. Both spatial mappings and correlation values show that repressive histone marks are less compartmentalized than active histone marks studied here, suggesting the different gene regulation mechanism of these two groups of histone marks. It is also interesting to note that DNA methylation instead of H3K27me3, is used upon gene repression at the late developmental stages of ES differentiation, suggesting a minor role of repressive histone marks in these cells^27^.

#### Replication timing

DNA replication occurs in a cell-type-specific temporal order known as the replication-timing (RT) program. RT reflects chromatin spatial organization in that early and late replication domains match well with compartments A and B, respectively^28^.

The degree of compartmentalization provides a more quantitative description of the 3D chromatin structure than the classification of a certain genomic region into compartments A and B. In Fig. 3a and 3b, we compare the degree of compartmentalization and replication timing. Strikingly, the degree of compartmentalization coincides very well with RT as can be seen from 3D structure (Fig. 3a) or one-dimensional (drawn as a function of the genome distance) profile (Fig. 3b). The values of Pearson correlation between degree of compartmentalization and replication timing is 0.85. Thus, it can be concluded that replication timing has a close relationship with local compactness. This modeling study thus also presents a vivid illustration of the relation between chromosome replication and structure.

### Spatial segregation between PMD and non-PMD

The widespread DNA hypomethylation in cancer methylome resembling the PMDs also occurs in IMR90 cell line^29^. The identification of PMD is presented in the SI text: PMD and non-PMD identification. By mapping the PMDs onto the chromatin structure, we found that PMDs are spatially segregated from non-PMDs (Fig. 3c). Nearly all of the PMDs (98.8% in length) locate in compartment B, suggesting the possible pathological change of this compartment during oncogenesis.

### Structural characterization of PMD and non-PMD

The contact probability along genomic distance derived from Hi-C experiments is significantly different between PMD and non-PMD (Fig. S9a and SI text: Comparison between the compactness of PMD and non-PMD). In contrast to non-PMD, PMD displays a much slower decrease in contact probability which means that PMDs are more compact than non-PMDs in the kilobase-megabase scale. We showed that this result is robust using Hi-C data from separate sources (Fig. S9b).

We further examined the Hi-C patterns for PMDs and non-PMDs longer than 300 kb. Consistent with the above contact probabilities, the Hi-C patterns for PMD and non-PMD show striking differences. For non-PMD, the majority possess a Hi-C pattern as shown in Fig. 4a which contains localized interaction domains and appears as several small squares along the diagonal of the Hi-C map. In contrast, the genomic regions inside each PMD contact nearly uniformly with each other and show a highly conserved pattern among PMDs typically as that shown in Fig. 4b.

**Figure 4.**
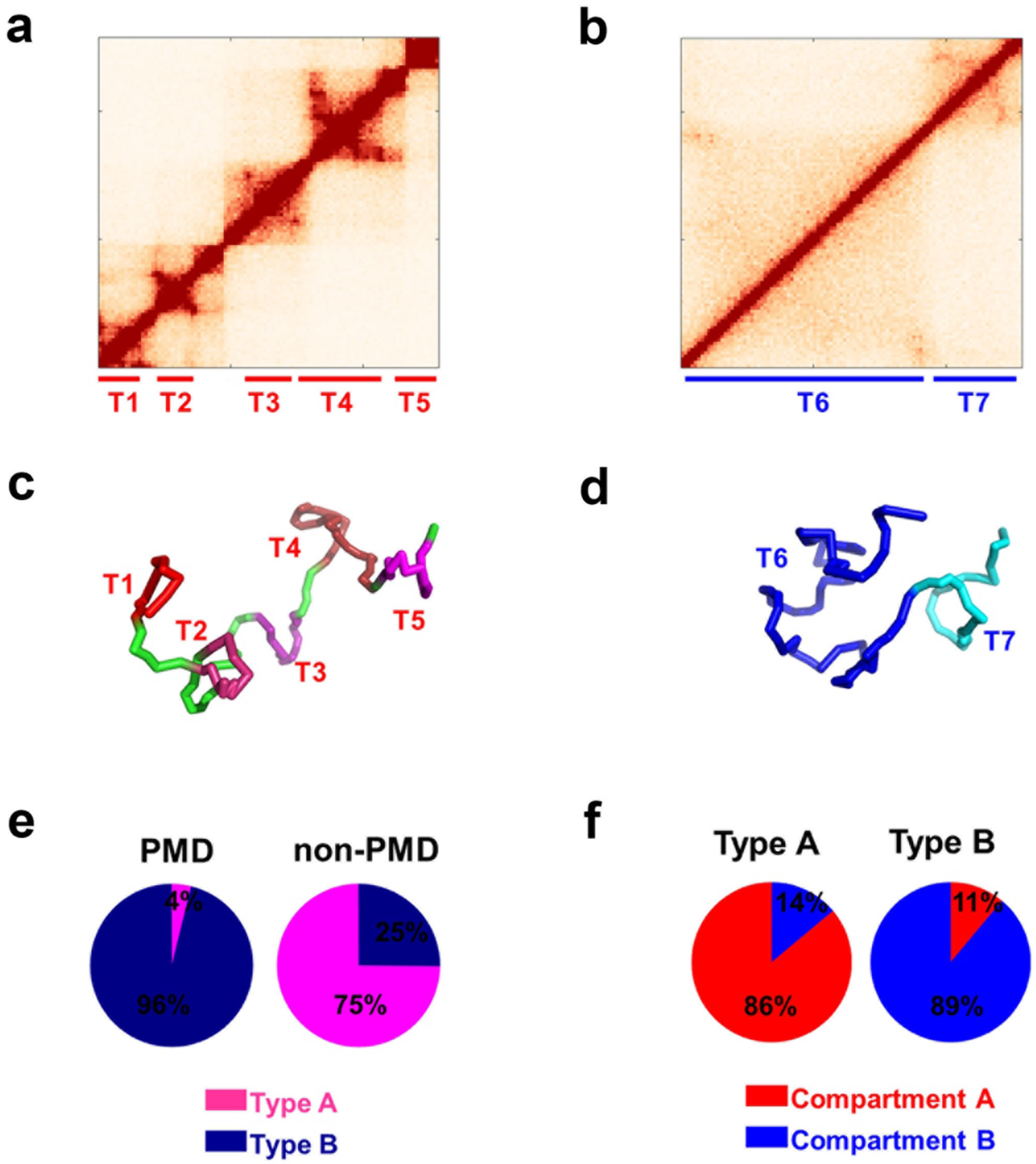
Characterization of type A and B chromatin. The Hi-C pattern of type A chromatin is different from type B chromatin: **(a)** Typical Hi-C of type A chromatin (chr1: 15.46-16.54 Mb). **(b)** Typical Hi-C of type B chromatin (chr1: 233.47-234.50 Mb). The structural modeling illustrates the different structure of type A/B chromatin: **(c)** Modeled structure of type A chromatin, corresponding to Hi-C data in (a). **(d)** Modeled structure of type B chromatin, corresponding to Hi-C data in (b). TADs are annotated in Hi-C data (a,b) and correspondingly colored in blocks in (c,d). **(e)** Composition of type A/B chromatin in PMD/non-PMD. **(f)** Composition of compartments A/B in type A/B chromatins.

To elucidate the structural differences between PMD and non-PMD, we construct 3D models for the two Hi-C patterns shown in Fig. 4a and 4b. The representative structures are presented in Figs. 4c and 4d, respectively. Considering the feasible computational cost for these segments of the chromatin, we used a 10-kb resolution of modeling at this domain scale in contrast to the 50-kb resolution in the modeling of the entire chromosome. Our modeling accurately captured the larger TAD numbers appeared in Fig. 4a than that in Fig. 4b. Similar to the comparison of compactness between compartments A and B, we can compare the local density or radius of gyration around the two modeled chromatin domains. Chromatin structure in Fig. 4a is obviously less compact than chromatin structure in Fig. 4b from our modeling (SI text: Estimation of chromosome spatial density).

### Clustering an individual chromosome into chromatin types A and B

As mentioned above, the typical Hi-C patterns for PMD and non-PMD are very different. Such an observation promoted us to further cluster all the PMDs and non-PMDs into two types, called here chromatin type A (typical as Fig. 4a) and chromatin type B (typical as Fig. 4b), according to their Hi-C patterns (see Methods). Nearly all the PMDs (96%) are clustered into type B and the ratio for non-PMD clustered into type A is 75% (Fig. 4e).

These two types of chromatin constitute 54.4% of the entire human genome (comparable to the 67.9% coverage of annotated TADs) and have an intimate relation with TADs and compartments. Types A and B chromatin are mainly located in compartments A and B, respectively (Fig. 4f). Considering that the majority of PMDs coincide with TADs (Fig. S8 and SI text: Comparison between the compactness of PMD and non-PMD), type A and B chromatins also share boundaries with TADs. Types A and B chromatin are complementary to TADs and can be correspondingly regarded as building blocks for compartments A and B.

The two types of chromatin domains are of similar average lengths (588 kb and 744 kb for types A and B chromatin, respectively). One type A contains on average 2.34 TADs, significantly more than that of type B (on average 0.99 TADs) (p-value < 10^-15^ by Welch’s unequal variances t-test). Therefore, the density of TADs also differs significantly between the two types of chromatin (3.99 TADs per Mb and 1.33 TADs per Mb for types A and B, respectively). The reason behind such large differences can be attributed to the different CTCF-binding site densities between types A and B which will be elaborated in the next section.

### Spatial segregation of chromatin loop according to loop size

Intrigued by the different Hi-C patterns and structures of chromatin types A and B, we interrogated their underlying sequence differences. They are found to be associated with GC rich and AT rich sequences, respectively. Most importantly, chromatin type A is 3.3 fold enriched for CTCF-binding sites (CBSs) than chromatin type B.

CBSs which are enriched at the boundaries of both TADs and chromatin loops play a crucial role in establishing chromosome structure^30^. The recently proposed loop-extrusion model also highlights the importance of CBS in that the locations of CBS alone can recapitulate most of Hi-C experimental results^14,15^. In the loop-extrusion model, cohesions acting as the loop-extruding factors, gradually form chromatin loops, but stall at TAD boundaries due to interactions with CTCF^14,15^. This model also unifies TADs and chromatin loops that TADs are formed by dynamic loops^15^. Thus in the following we regard TAD and chromatin loop as exchangeable.

We found that the density of CBS in compartments A and B also differs significantly (26.6 per Mb and 12.4 per Mb in compartments A and B, respectively). Consistent with the loop-extrusion model^14,15^, the higher CBS density in chromatin compartment A could result in smaller chromatin loops compared to those in compartment B, which is in fact suggested by the different Hi-C patterns of the two chromatin types. The contrast of GC-content and CBS density in A/B compartments can also be easily seen by mapping these properties onto the chromatin model (Fig. 5a and 5b).

**Figure 5.**
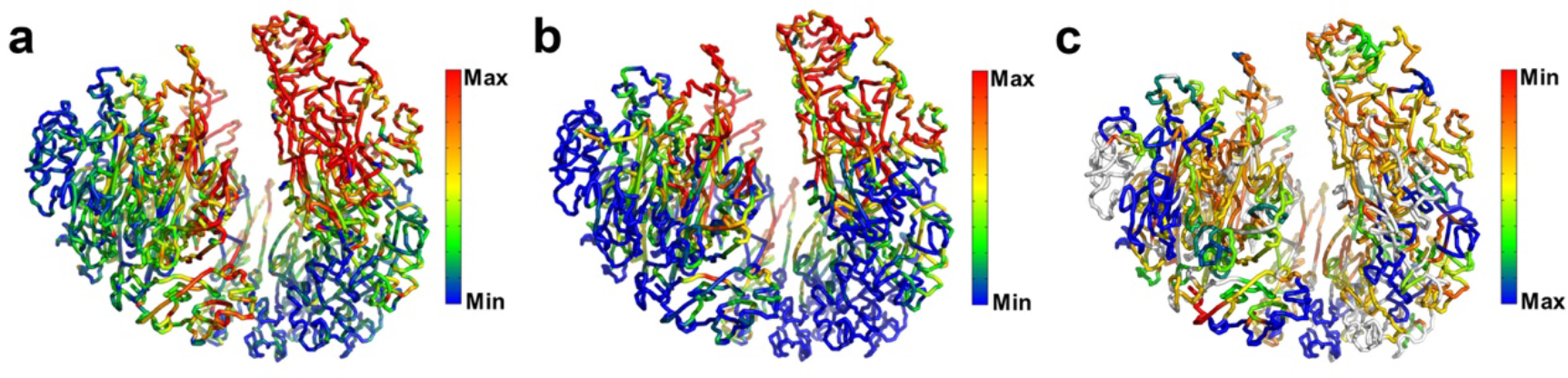
Mapping sequence properties and TAD size onto the 3D chromatin structure. **(a)** CpG density (Colored as the count of CpG in a 50-kb chromosome region). **(b)** Density of CTCF binding site (Colored as the count of CTCF binding site in a 400-kb chromosome region). **(c)** TAD size (Colored as the size of TAD in which the genomic region belongs). Compared to chromatin compartment B, compartment A is enriched with CpG dinucleotide, CTCF binding site and has smaller TAD.

The spatial segregation between the two compartments implies the chromatin loop segregation according to their sizes, i.e., large loops tend to assemble with each other near nuclear periphery and small loops also spatially accumulate in the nuclear center. The spatial segregation of loop size (or TAD size) can also be easily seen from our mapping of TAD size onto the chromatin model (Fig. 5c).

## Discussion

We observed the spatial segregation of chromatin loops according to loop sizes. Such a phenomenon strongly suggests the role of entropy in chromatin organization due to “depletion attraction”^31,32^.

The depletion attraction only occurs in crowded environment. Consider many large and small spheres confined in a space. When two large spheres contact with each other, their excluded volumes overlap which leads to the increase of available volume for the small sphere. Thus, the entropy of small spheres and then the system entropy paradoxically increase. The large sphere also tends to contact with the boundary of the confined space due to depletion attraction. A schematic of the excluded volume or entropic effect can also be seen from Fig. 1A in ref 33. Given the highly crowding cellular environment, such non-specific entropy force might contribute to many processes in cells^32,34,35^. Another elegant example is that the replicated bacteria daughter strands spontaneously segregate due to entropic forces^36^.

Aggregation of large chromatin loops causes the overlap of excluded volumes surrounding them, and increases the accessible volume and entropy for small chromatin loops. As a result, the segregation of chromatin loops according to their sizes could increase the system entropy. Here, we propose that an entropy-driven mechanism can explain the chromatin compartmentalization. The segregated small (large) chromatin loops comprise chromatin compartment A (B). The nuclear envelop can be regarded as a large entity which tends to aggregate with large chromatin loops. Thus, chromatin compartment B with larger chromatin loops than compartment A prefers to locate around the nuclear membrane.

Next, we argue that chromosomes positioning in the nucleus also correlates with chromatin loop size. The physical mechanisms underlying chromosome positioning remain largely unknown. Previous studies have shown that chromosomes position in the nucleus in an activity-based way^37^. Gene-rich chromosomes are situated at the center of the nucleus, at the same time the more gene-poor chromosomes concentrate towards the nuclear periphery^37^. It should be noted that the proteins of inner nuclear membrane may be not necessary for localizing chromosomes at the nucleus periphery^37^, suggesting the existence of non-specific driving forces.

In addition, the positions of several chromosomes are well determined in FISH experiments. Gene-rich chromosomes 1, 16, 17, 19, 22 are close to the nuclear center while gene-poor chromosomes 2, 13, 18 are close to the nuclear periphery for fibroblast cell^37^. For these chromosomes, we calculated the CBS density and the median size of chromatin loops identified in Hi-C experiments (Table 1). We found that gene-rich chromosomes have much higher CBS densities and are expected to have smaller chromatin loops than gene-poor chromosomes according to the loop-extrusion model. This is indeed the case as can be seen from the median size of chromatin loops identified from experimental Hi-C data (Table 1).

**Table 1.** Density of CTCF-binding site and median size of chromatin loops in different chromosomes.

| **Chromosome** | **CBS density (Mb<sup>-1</sup>)** | **Loop median size (kb)** |
| --- | --- | --- |
| 1 | 17.5 | 200 |
| 16 | 16.7 | 180 |
| 17 | 26.6 | 150 |
| 19 | 30.1 | 110 |
| 22 | 17.9 | 160 |
| 2 | 14.6 | 240 |
| 13 | 9.7 | 303 |
| 18 | 12.0 | 240 |

Thus, the median loop size of chromosome correlates well with chromosome positioning: chromosomes with smaller median loop size segregate to the center of nucleus and larger median loop size to nuclear periphery. Particularly, chromosomes 18 and 19 which have similar sizes show strikingly different CBS density and chromosome position.

Therefore, the size of chromatin loop plays an important role in determining chromatin structure at both intra- and inter-chromosome levels. An individual chromosome segregates into two compartments and chromosomes positions in the nucleus, both according to chromatin loop size.

We summarize the chromosome folding principle proposed here in Fig. 6: The CG-rich DNA sequences have higher densities of CTCF-binding site than AT-rich DNA sequences. Consistent with the loop-extrusion model^14,15^, the CG-rich DNA sequences have smaller chromatin loops (or TADs). Chromatin loops segregate according to their sizes both within a single chromosome and among the whole chromosomes, suggesting the excluded volume or entropic effect in genome spatial organization. Within a single chromosome, the larger chromatin loops tend to aggregate around the nuclear boundary, leading to chromatin compartmentalization. Among the different chromosomes, the ones having larger chromatin loops prefer nuclear periphery resulting in chromosome positioning.

**Figure 6.**
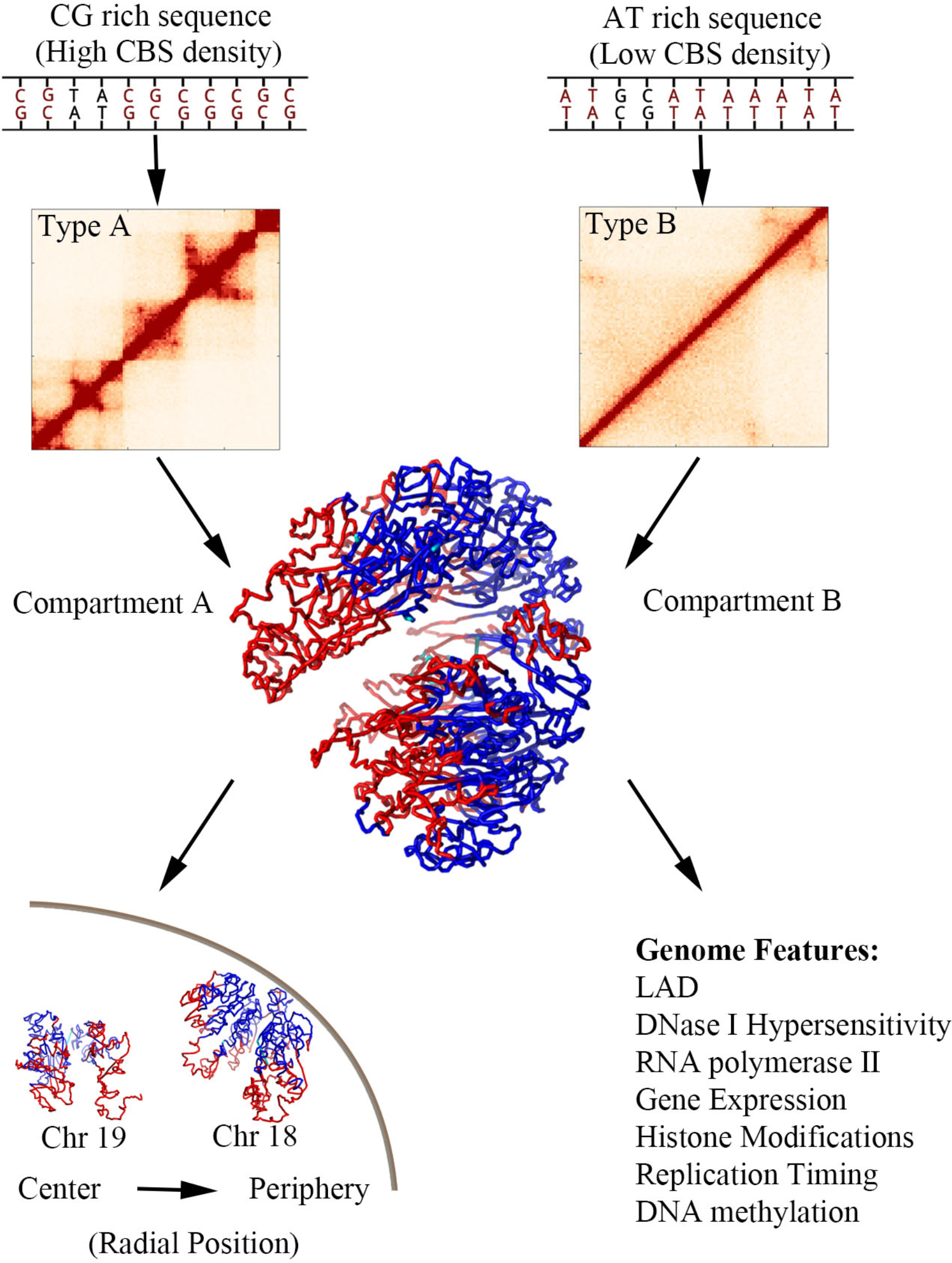
Schematic of the folding principle of chromosome. Chromatin type A is enriched with CG sequences and CTCF-binding sites. Thus, chromatin type A has relatively smaller chromatin loops compared to chromatin type B. Larger chromatin loops tend to aggregate around nuclear membrane, suggesting an entropy-driven chromatin compartmentalization. Chromosomes with larger median chromatin loops prefer nuclear periphery, also implying an entropy-driven chromosome positioning. The distribution of various genome features are related to chromatin compartmentalization.

## Conclusions

Using Hi-C experimental data and a restraint-based modeling strategy, we built chromatin spatial structures. The correlated spatial segregation of compartments and various genome features was observed. Previous studies mainly focused on the colocalization of genomic features on a linear map. Here the spatial mapping provides a simple relation between various genome features. Their co-segregation in the 3D chromatin structure results in patched distributions and colocalization along the genome. We also argue that the spatial segregation of chromatin loop according to loop sizes strongly suggests the entropic effect in chromatin compartmentalization and chromosome positioning.

## Methods

### Structural modeling

The structural modeling process for chromosome is mainly composed of two procedures: (1) converting Hi-C contact frequencies to distance restraints used in the modeling and (2) constructing and optimizing the structure of the coarse-grained chromosome model to satisfy the above distance restraints. Details are described below.

1. Converting Hi-C contact frequencies to distance restraints

The spatial distance *d* between two genomic segments scales with genomic distance *l* is written as 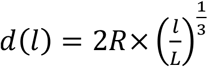 where *L* is the total genomic length of all human chromosomes (set as 6 Gb) and *R* is the radius of cell nucleus (set as 10 μm). The 1/3 scaling is in accordance with the 3D-FISH experimental results^38^. The Hi-C experimental data provide the relation between the contact frequency *C(l)* and the genomic distance *l*. Contact frequency can therefore be regarded as a function of spatial distance, which is also the underlying foundation of structural interpretation of Hi-C data^13^. We then mapped contact frequencies to spatial distances by linear interpolation on the double-logarithm plot of *C(l)* and *d(l)*.

Mathematically, the above mapping strategy can convert all the contact frequencies into restraint distances. However, for two genomic segments that are far from each other in the genomic sequence, their contact frequency is generally very low and the corresponding Hi-C data have low signal/noise ratios, which may introduce large errors in the estimation of the restraint distance. To alleviate the influence of such artifacts, contact frequencies with genomic distances larger than 2-Mb are smoothed by constructing (200kb×200kb)-blocks in these regions of the contact matrix. The contact frequency averaged over a block was then assigned to each segment pair inside it.

1. Optimizing the structure of the coarse-grained chromosome model

We coarse-grained a chromosome as a string of beads, each bead representing one genomic segment. The equilibrium distance between two consecutive beads was also obtained by mapping the contact frequency to the spatial distance as mentioned above. A set of randomly generated initial structures were used for further structure optimization using MD simulations.

Two types of potentials are used in our method: a bond-stretching term 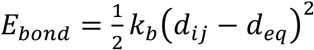 applied to consecutive beads to maintain the continuity of the chromosome polymer chain, and a restraint potential 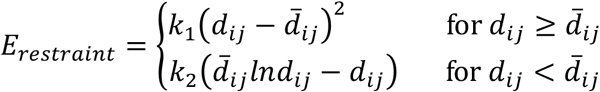 to force the positions of beads to follow the distance restraints obtained from Hi-C data, where *d_ij_* is the simulated spatial distance between beads *i* and *j, d_eq_* is the equilibrium distance between two neighboring beads, *k_b_* is a parameter for the harmonic potential, *k*_1_ and *k*_2_ are set according to restraint distances so that the reproduction of Hi-C contact at shorter restraint distances is of higher priority.

In order to avoid too large movements in each step of the MD simulation and achieve a high computational efficiency at the same time, the time integration interval was set to be inversely proportional to the maximum force applied on all the beads. Velocities were randomly generated every 100 steps based on the Boltzmann distribution to accelerate the conformation search. We ran each simulation for 80,000 steps, by when the conformation of polymer chain would change little, suggesting that a stable chromosome structure has been obtained (Fig. S1a). The final conformations optimized from different initial structures were collected for further statistical analysis. We generated 300 models for chromosome 1 of IMR90.

With the modeled chromatin conformation ensemble, we can generate the simulated Hi-C contact matrix by an inverse mapping, from spatial distances to Hi-C contact frequencies. For each conformation, we calculated the spatial distances between all the bead pairs which were then converted to contact frequencies. The modeled contact matrix was obtained by an average over the 300 simulated models.

### Clustering for chromatin types A and B

We re-classified PMDs and non-PMDs of all chromosomes into two types according to their Hi-C patterns. As we have observed, the variances of contact frequencies along the genomic distances differ significantly between typical PMDs and non-PMDs. Therefore we chose contact variances (in logarithmic scale) at different genomic distances (in logarithmic scale) as the classification feature. We first used long segments to train and validate the linear discriminant analysis (LDA) classification model, then we applied the trained model on all segments to classify them into structurally meaningful types.

All segments longer than 500-kb were first clustered into two sets using k-means. While 68% of long non-PMDs were grouped into the first set, almost all the long PMDs (97%, 593 out of 613 segments) were grouped into the second. We thus labeled the first set as segment type A and the second as type B. All these long segments were then randomly divided into a training set (80%, 850 segments) and a test set (20%, 213 segments). To avoid over-fitting, we trained the LDA classifier on the former set with 5-fold cross validation. The cross validation gave a result of only 0.59% loss, demonstrating an almost perfect match of our model to our training set. By further applying the model to the independent test set, we successfully predicted the structural types of the test segments with an error rate of 1.9%, only 4 out of 213 test data gave wrong prediction.

Using this validated classifier, we grouped all the 2508 PMD and non-PMD segments into two types. While 75% (1116 out of 1491) and 25% (375 out of 1491) non-PMDs belong to type A and B respectively, almost all the PMDs (96%, 977 out of 1077) belong to type B.

The source code for structural modeling and segment classification are available at http://www.chem.pku.edu.cn/gaoyq/Chromatin/codes.tar. The Hi-C patterns of PMD and non-PMD are also available at http://www.chem.pku.edu.cn/gaoyq/Chromatin/HiC_PMD_nonPMD.zip.

## Statistics

Welch’s unequal variances t-test was used to calculate the statistical significance of the TAD number difference between type A and B.

## Acknowledgements

The work was supported by National Natural Science Foundation of China (21233002, 21573006 and 91427304). We would like to thank Prof. Fuchou Tang, Dr. Fan Guo, Lin Li and Rui Zhou for helpful discussion.

## Authors’ contributions

W.J.X. and Y.Q.G. conceived of and designed the study. W.J.X. led data analyses. L.M. performed chromatin modeling and structural analyses. S.L. processed experimental data and performed Hi-C clustering. L.Z. and X.C. contributed to data analyses. Y.Q.G. supervised the study. W.J.X. wrote the manuscript with the inputs from L.M. and S.L. All authors contributed to the final version of the manuscript.

